# Fast and Accurate Estimation of Gas-Phase Entropy from the Molecular Surface Curvature

**DOI:** 10.1101/2021.05.26.445640

**Authors:** Vishwesh Venkatraman, Amitava Roy

**Affiliations:** NTNU, Department of Chemistry, 7491 Trondheim, Norway; Bioinformatics and Computational Biosciences Branch, Office of Cyber Infrastructure and Computational Biology, National Institute of Allergy and Infectious Diseases, National Institutes of Health, Hamilton, MT 59840, USA; Department of Computer Science, University of Montana, Missoula, MT, USA

## Abstract

Estimating entropy is crucial for understanding and modifying biological systems, such as protein-ligand binding. Current computational methods to estimate entropy require extensive, or at times prohibitively extensive, computational resources. This article presents SHAPE (SHape-based Accurate Predictor of Entropy), a new method that estimates the gas-phase entropy of small molecules purely from their surface geometry. The gas-phase entropy of small molecules can be computed in ≈0.01 CPU hours with run time complexity of 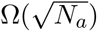, where *N*_*a*_ is the number of atoms. The accuracy of SHAPE is within 1 − 2% of computationally expensive quantum mechanical or molecular mechanical calculations. We further show that the inclusion of gas-phase entropy, estimated using SHAPE, improves the rank-order correlation between binding affinity and binding score from 0.18 to 0.40. The speed and accuracy of SHAPE make it well-suited for inclusion in molecular docking or QSAR (quantitative structure-activity relationships) methods.

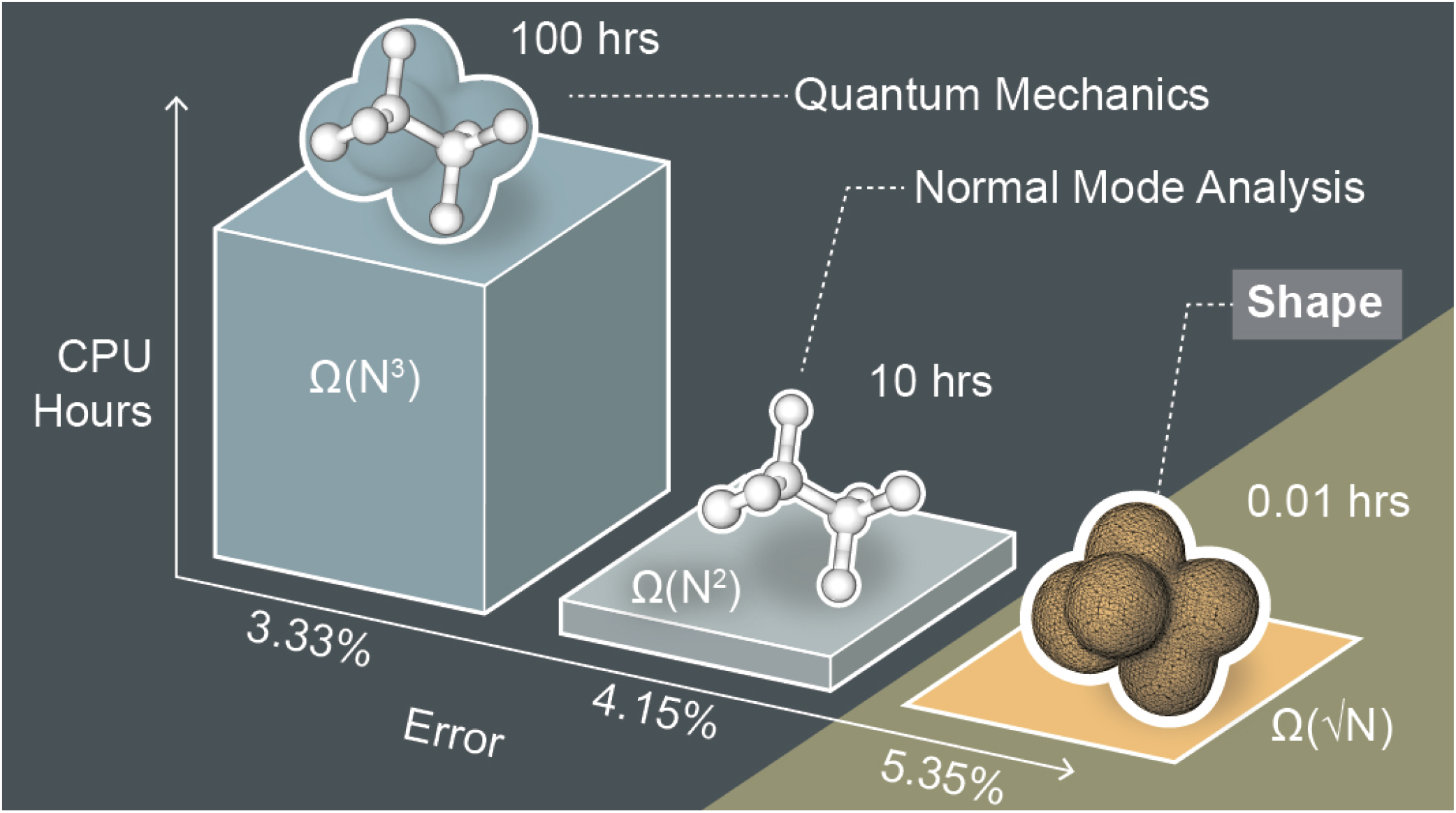

## Introduction

Free energy governs all chemical processes. Change in free energy in a chemical process can be written as Δ*G* = Δ*H* −*T* Δ*S*, where *G* is the Gibbs free energy, *H* is the enthalpy, *S* is the entropy, and T is the absolute temperature. The enthalpy represents interactions between atoms, e.g., hydrogen bonds, electrostatics, steric clashes, and can be estimated efficiently with computational techniques. The entropy in a chemical process represents the relative probability of the molecules being in a particular state, called a microstate, and is challenging to estimate computationally. Entropy depends on the dynamic flexibility of the system. All possible microstates that a molecule can exist in need to be calculated to determine the probability of a single microstate. Experimentally, the change in total entropy in a chemical process is measured by isothermal titration calorimetry. Conceptually, in computational exercises, entropy is divided into different terms based on the types of dynamics a molecule exhibits, e.g., rotational, translational, vibrational, conformational. If we want to understand and manipulate a biological system, such as protein–ligand binding, we need an efficient estimation of the change in entropy in a chemical process. Computational methods for estimating gas-phase entropy are currently time-consuming and resource-intensive. Here, we present a fast and accurate method for estimating the gas-phase entropy of small molecules, with computation requiring only CPU seconds, and with a small (1 − 2%) loss in accuracy.

### Entropy in docking

Virtual screening has become a key component to understand protein-ligand binding in drug discovery pipelines, and molecular docking is an essential part of such pipelines. All molecular docking methods use a docking score to rank the docked compounds. Currently, these methods can predict the pose of a ligand when both the binding pocket and the ligand are known.^1,2^ However, docking programs perform poorly while predicting the ligand binding affinity ranking or when attempting to classify active and inactive compounds from binding affinity prediction.^1,2^ Part of the difficulty lies in the lack of a suitable representation of the binding entropy in the scoring functions.^3–6^ When an entropic contribution is included in scoring functions, it is assumed that the significant entropic contribution to binding comes from the conformational changes of the protein and ligand; however, this assumption is not well-founded.^5,6^ The question remains whether the flexibility of a subset of sidechains is sufficient to capture entropic changes in a binding event.^7^ Indeed, an increasing collection of computational and experimental studies indicate that the vibrational pattern of the protein and the ligand can contribute to changes in the binding entropy.^6,8^ Moreover, when the vibrational entropy is the dominant factor in a binding event, the effect may come from all vibrational frequencies rather than a select few.^6,9^

For computational purposes, the gas-phase entropy can be divided into translational, rotational, and vibrational entropy terms. In practice, these terms are calculated from the most probable conformation of the molecule. The difference between the gas-phase entropy calculated from a single conformation and those averaged over multiple possible conformations is the conformational entropy. Calculation of the gas-phase entropy, especially full vibrational spectra and the corresponding vibrational entropy, is prohibitively time and resource-consuming for high-throughput docking. For example, a density functional theory (DFT) based quantum mechanical calculation of the vibration spectrum takes hundreds of CPU hours, while a normal mode analysis (NMA) using molecular mechanics takes CPU seconds, but the underlying parameterization for the force field takes tens of CPU hours for each ligand. The runtime complexity of quantum mechanical calculation of vibrational entropy depends on the complexity of the underlying theory. For complicated systems, it can be an NP-hard problem. In our experience, the runtime complexity of quantum mechanical calculation of vibrational entropy is at least cubic 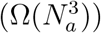, and that of NMA is quadratic 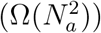, where *N*_*a*_ is the number of atoms.

This article describes a fast and accurate method to estimate molecular gas-phase entropy, called SHAPE (SHape-based Accurate Predictor of Entropy). SHAPE does not require additional parameterization for a ligand, takes CPU seconds for calculation, and has a square root runtime complexity of 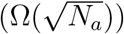. SHAPE uses an informational theory-based approach to calculate the entropy, and its accuracy is only 1-2% lower than DFT or NMA based calculations. Both the accuracy and speed of computation of SHAPE make it an excellent candidate for scoring functions in high-throughput docking.

### Entropy in thermodynamics and informational theory

Information entropy, first introduced by Shannon,^10^ is a measure to gauge the uncertainty, or lack of information, associated with a distribution function *H* of *n* probable outcomes, with probabilities *p*_*i*_,

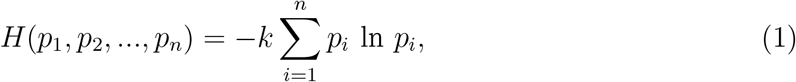

where ∑_*i*_ *p*_*i*_ = 1 and *k* is a positive constant. Shannon entropy is a statistical idea and may not have a physical meaning. For a physical system, if microscopic states arising from a macroscopic physical constraint can be identified, its entropic energy can be calculated using Gibbs entropy, which takes the form of Equation 1 with a specific constant, Boltzmann constant (*k*_*B*_). In such a case, the probability *p*_*i*_ is the probability of the microstate *i*, given the macroscopic physical constraint. Gibbs entropy is also referred to as Gibbs-Shannon entropy due to the similarity of the equation.

Entropy can also be expressed as a function of the number of ways the atoms or molecules in a thermodynamics system can be arranged (microstates) under a macrostate’s constraints, as expressed by famous Boltzmann’s equation:

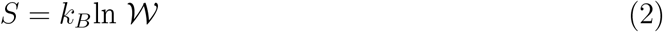

where *S* is entropy, *k*_*B*_ is Boltzmann’s constant, and 𝒲 is the number of microstates corresponding to the macrostate. The Gibbs-Shannon entropy is a more generalized formulation of the Boltzmann entropy and equals the Boltzmann entropy only under certain equilibrium conditions.^11^

The change of entropy in athermodynamic process, such as binding a protein and a ligand, can be calculated using the Gibbs-Shannon or Boltzmann formulation if the relevant microstates associated with the event can be identified and their probabilities can be calculated. For protein-ligand binding, macroscopic constraints can be the temperature, presence of a solvent, protein, ligand, and protein-ligand complex concentration. Microscopic states can be rotational, translational movements of the full molecules, vibration, flexibility patterns of different atoms, and the orientation of solvent molecules.

### Estimation of gas-phase entropy

Shannon’s entropy has been applied in density functional theory (DFT) to calculate chemical properties, a review of related works can be found in Rong et al. ^12^. For example, it has been applied to electron density topological functions, such as the Fukui function, to calculate molecular ionization.^13^ When applied to the molecular electron distribution, it can identify the delocalization of electrons or the lack thereof. This article applies Shannon entropy to molecular shape functions derived from van der Waals radii to calculate gas-phase entropies of small molecules. To the best of our knowledge, this is the first application of Shannon entropy to the surface curvature to estimate gas-phase entropy.

The estimation of gas-phase entropies has largely relied on *ab initio* molecular orbital theory.^14–16^ Density functional theory with different functionals and basis sets has been frequently employed in these calculations.^17,18^ Other methods based on semi-empirical orbital theory^15,19^ and molecular dynamics simulations^20–22^ have also been used to estimate entropies. Many of these methods however, require considerable time and computational resources, and are therefore not suitable for applications where millions or billions of such calculations are required, such as in virtual screening for drug discovery. Group additivity methods,^23–25^ on the other hand, are fast and work by dividing the molecule into distinct groups and estimating the entropy based on the individual group parameters. The efficacy of the method is however limited by the fact that the list of groups cover only a limited range of chemistries.

To date, the gas-phase entropy has been estimated from the dynamics and distribution of molecular chemical properties; however, it has never been estimated (to the best of our knowledge) from the shape curvature. This article shows that the gas-phase entropy can be calculated from the curvature of the molecular surface alone. We calculated the Shannon shape entropy for a set of 1212 organic molecules from their curvatures estimated from the electron isodensity surfaces and approximated by the van der Waaals (VDW) radii. In both cases, the Shannon shape entropy was found to have a linear relationship with the experimentally determined gas-phase entropies with a coefficient of determination (*R*^2^) of ≈0.90. We compared the performance of Shannon shape entropy, calculated from the VDW surface against gas-phase entropies calculated using quantum-chemical Gaussian-4 (G4) level of theory and normal mode analysis (NMA). In comparison with G4 or NMA methods, we show that SHAPE speeds up the entropy calculation by orders of magnitude while losing only 1-2% accuracy (representing ≈0.6 Kcal/mol energy at room temperature).

### Inclusion of gas-phase entropy in docking score

To investigate the effect of including gas-phase entropy in the docking score, we calculated the binding affinity of 22 thrombin inhibitors using the SMINA^26^ scoring function with and without the gas-phase entropy as a proof of concept. The inclusion of gas-phase entropy in the scoring function showed trends of improvement in the *R*^2^ values from 0.23 to 0.40; for Kendall’s *τ*, a rank correlation coefficient, values were improved from 0.18 to 0.40. The authors acknowledge that for a proper treatment of entropy in the docking score, one should include the entropy change in protein and ligand upon binding, rather than the entropy of the ligand in the bound conformation. However, in our dataset, all the 22 ligands bind to the same pocket; consequently, the differences in gas-phase entropy between the ligands is a first-order approximation of the differences in binding entropy of the corresponding protein-ligand complexes. This article presents the power of shape entropy in estimating thermodynamic entropy. A full treatment of binding entropy should include contributions from the protein and the solvent and is a work in progress.

## Methods

### Data Curation

As the first step, we built a database with experimental gas-phase entropy values (at 25*°*C in *J* /*mol* · *K*) for a various organic compounds (involving elements C,H,N,O,S,P,Cl,Br and I) curated from literature.^14,16,27,27,28^ Overall, 1212 compounds with corresponding experimental entropies in the range 180 – 855 *J* /*mol*·*K* (≈13-61 kCal/mol at 300K) were obtained (see Figure 1(E)). A significant majority of the compounds had entropies below 500 *J* /*mol* · *K* and had less than five rotatable bonds. High entropies (*>* 700 *J* /*mol* · *K*) were associated with compounds containing more than 45 heavy atoms and more than ten rotatable bonds. We also curated vibrational entropy calculated by quantum-chemical Gaussian-4 (G4) level of theory and by normal mode analysis (NMA) of 1091 and 389 of those 1212 compounds, respectively.^27^ Rotational and translational entropies for those molecules were calculated analytically from their shapes to determine the remaining terms.^27^ In our curation of data calculated by the G4 or NMA method, only the most probable conformation of the molecules was considered. Consequently, those values do not include a contribution from the conformational entropy. However, a recent study identifies that conformational entropy accounts for ≈*<* 5% of gas-phase entropy in small molecules.^29^ To minimize the error of non-inclusion of conformational entropy in the G4 and NMA calculation, we fitted the calculated data to the experimental values before calculating any statistical properties of those values.

**Figure 1:**
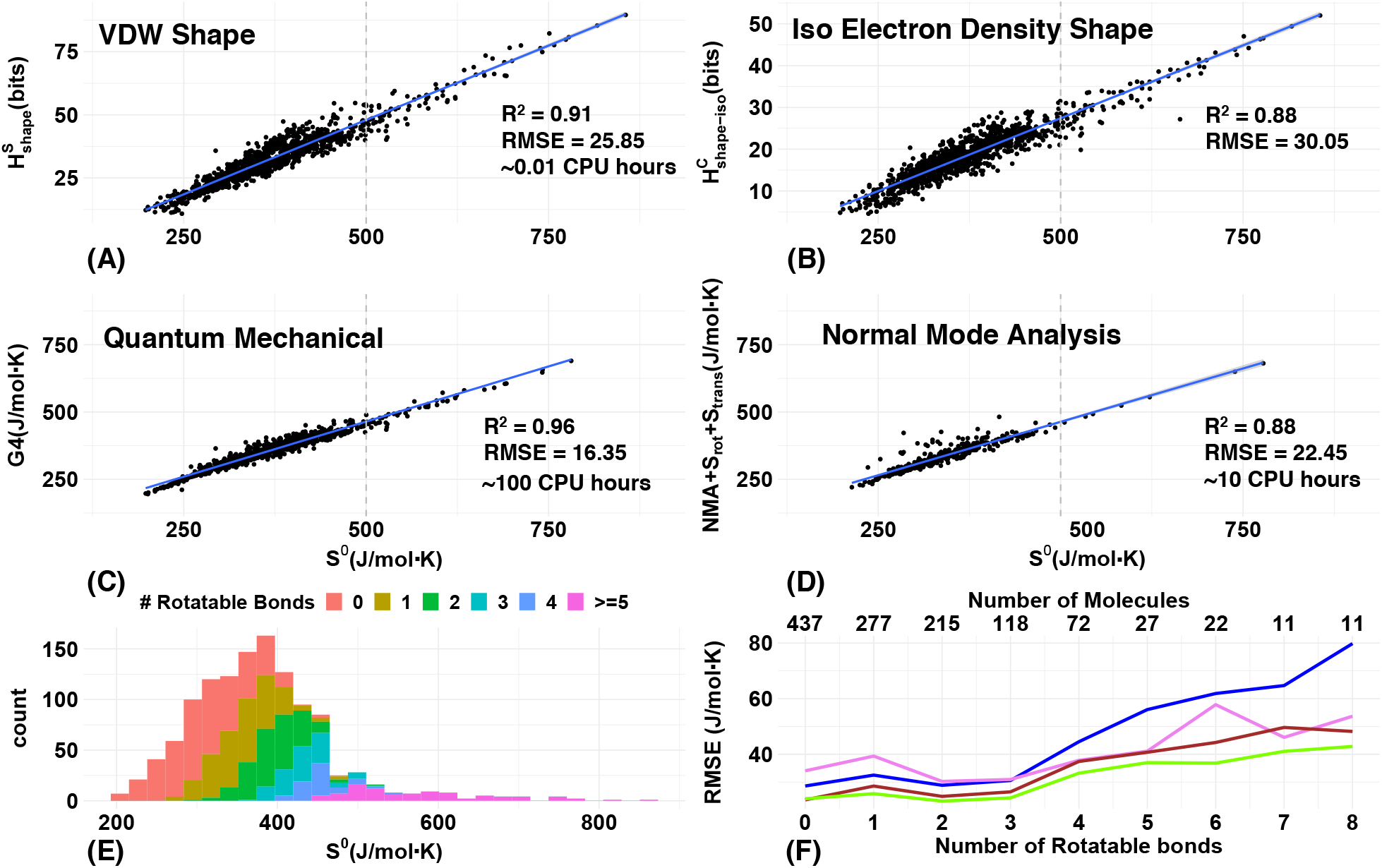
Comparison of the experimental gas-phase entropies and shape entropies computed using the shape index of VDW surface without hydrogens and curvedness of isodensity surface of electrons with hydrogens are shown in plots (A) and (B). Both (A) and (B) show co-linearity between shape and gas-phase entropy. Plot (C) compares experimental gas-phase entropies and calculated gas-phase entropies using G4 quantum chemical calculations, and plot (D) compares experimental values with gas-phase entropy calculated by normal mode analysis (NMA). For both G4 and NMA calculations, translational and rotational entropies were calculated analytically from the molecules’ shape. Plot (E) shows the histogram of the experimental entropies. Plot (F) shows the root mean square error (RMSE) from the linear fitting of *H*_*shape*_ to *S*_*gasphase*_ as a function of the number of rotatable bonds of the molecules (x-axis). The number of molecules with a specific number of rotatable bonds is shown in the X2 axis. To account for the flexibility of the molecules, we generated up to 10 conformation clusters, identified the cluster representatives, calculated the heat of formation to build the representative conformations, and averaged the Boltzmann weighted *H*_*shape*_ values of the representative conformations to calculate the predicted *S*_*gasphase*_. Blue and violet lines show RMSE in *H*_*shape*_ calculated from curvedness 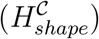 and shape index 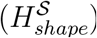, respectively. Brown and green lines correspond to the RMSE from Boltzmann-weighted 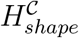 and 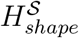, respectively. Boltzmann weighting reduces the RMSE in 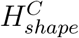 and stays approximately the same for 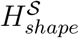. Note that calculation of vibrational entropy by G4 method takes 100s of CPU hours. The same takes about CPU seconds with NMA; however, the NMA method with molecular mechanics force field requires 10s of CPU hours per ligands for parameterization. Entropy can be calculated using CPU seconds of resources with SHAPE, the shape-based entropy method, and does not need additional parameterization for new ligands. SHAPE is an ideal candidate for the docking score in high throughput docking exercises.

### Molecular structure calculation

Our curated database contains SMILES strings of the chemical compounds. The SMILES strings for each molecule were converted into a single set of 3D coordinates using RDKit.^30^ The van der Waals (VDW) radii of the atoms were assigned using the software OpenBabel.^31^ Note that the VDW assignment by OpenBabel does not depend on the local environment. For example, all carbon atoms will have the same VDW radius, and no further parameterization is needed for individual small molecules. The structures were subsequently minimized using the Universal Force Field^32^ (UFF) implemented in RDKit. Similar to the VDW parameters, apriori parameterization of the charges is sufficient and any additional parameterization is not required.

### Molecular surface calculation

Once a molecular structure was calculated, the molecular surfaces were calculated from those structures by two different methods.

#### i) VDW surface calculation

In this study, an analytical form previously studied by Gabdoulline and Wade ^33^ has been used to calculate the surface and is defined as *M* = *G*^*−*1^(*c*) where the function *G* is a weighted sum of atom-centered Gaussians given by:

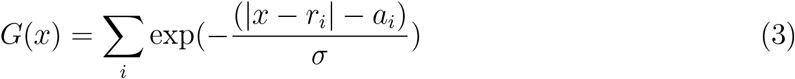

where *r*_*i*_ is the position of the *i*^*th*^ atom centre and *a*_*i*_ the corresponding VDW radius. The smoothing parameter 0 ≤ *σ* ≤ 1 affects the level of detail associated with the surface. Larger values of *σ* smooth out the details of the surface, while the smaller values of *σ* preserve more details of the surface feature. Please see the *SI* (Table ST1) to compare surface details at different *σ* values. The Gaussian surface isovalue *c* controls the volume enclosed by the surface. In this article, we calculated the surfaces at the isovalue *c* = 1.0, and used a smoothing factor of 0.1.

#### ii) Electron density surface

Isodensity surfaces are defined as surfaces around a molecule at which the electron density has a constant value. We calculated isodensity surfaces at the isodensity level 0.007 *e*^*−*^Å^*−*3^ for the molecules using the program ParaSurf^34^ that uses the output from a semi-empirical molecular orbital program such as MOPAC.^35^

### Surface curvature

The VDW surface for the molecules defined by Equation 3 is a continuous differentiable function and defined analytically at every point. From the analytical expression of the surface, we can calculate principal curvatures at every point. If we can draw a small normal plane at a point on a surface, and calculate the curvature of every line going through the point, then the highest and lowest curvatures of those lines, calculated at the point, are the principal curvatures *κ*_1_ and *κ*_2_ (*κ*_1_ *> κ*_2_) of the surface at that point. The principal curvatures measure how the surface bends in different directions at a point (see do Carmo^36^) and can be calculated analytically from the first- and second-order partial derivatives of the surface function.^37,38^ The principal curvatures can be further combined to define the curvedness (𝒞) and shape index (𝒮):^39^

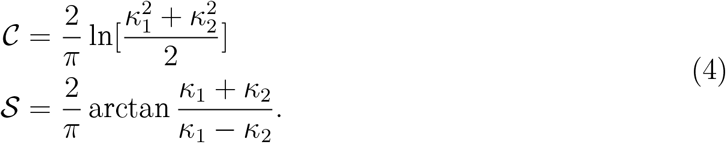

Both 𝒞 and 𝒮 measure the local shape of a surface. While 𝒞 can vary between −∞ and ∞, 𝒮 varies from -1 (concave) to 1 (convex).

In the absence of an analytical form for an iso-density surface calculated from an electron density, the curvedness and shape index values were calculated numerically. The output of the ParaSurf surface files was first converted to .obj geometry definition file format, proceeding which, curvature calculations were carried out using routines in the Visualization and Computer Graphics Library.^40^

### Shannon shape entropy

The molecular surface, once calculated, is discretized by a triangle mesh using the GNU triangulated surface library,^41^ i.e., covering the surface with triangles. After triangulation, the curvedness and shape index values are calculated at the center of each triangle of the mesh. Given *ntri* triangles in the surface mesh, the Shannon shape entropy can be written as:

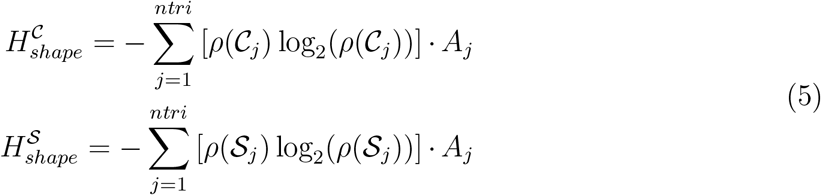

where the *A*_*j*_ is the area of the *j*^*th*^triangle and *ρ*(𝒞_*j*_) and *ρ*(𝒮_*j*_) are the probabilities of having of a curvedness value 𝒞_*j*_ and shape index value 𝒮_*j*_, respectively. The probabilities are determined by binning the curvedness and shape index values on a molecular surface. The calculation of the entropy using equation 5 can be impacted by the bin-width as well as the smoothing factor of the Gaussian surface function. We varied the bin numbers between 64, 128, and 256 and the smoothing factor *σ* between 0.1, 0.3, and 0.5. We then calculated the Pearson correlation coefficient between the shape entropies and the experimental gas-phase entropy to determine the effect of *σ* and the number of bins (see *SI*, Figure SF1). While the smoothing factor *σ* has a strong effect on the correlations, the number bins have only weak influence on the values (see *SI*, Figure SF1). We chose 64 bins to calculate the histograms and the probability density of the curvedness values using *σ* = 0.1 for the rest of the work.

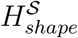 is mostly sensitive to major variations of local shapes. For example, we measured the 𝒞 and 𝒮 for a carbon atom. The VDW surface of a carbon atom is spherical, and we expect the *H*_*shape*_ to be zero. Values of 𝒞 show slight variations from the surface due to the artifacts from triangulation; consequently, 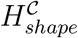 is a small non-zero number. However, 𝒮 is less sensitive to the triangulation effect, and 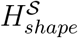 is zero.

The gas-phase entropy of molecules was modeled from the Shannon shape entropy as:

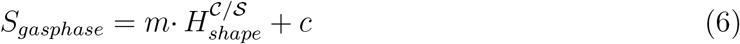

where *m* and *c* are constants and determined from fitting the *H*_*shape*_ to experimental data.

### Boltzmann weighted average

To account for multiple possible conformations of molecules, we generated conformers using RDKIT. Starting from the SMILES representation of a molecule, up to ten conformers were generated using RDKIT. Since the generated conformers can be structurally similar to each other, only conformations that are at least 0.5Å root mean square deviation (RMSD) apart from one another were retained. As a result of this filtering, the final tally of generated conformers may at times be less than 10. Each conformation was then subjected geometry optimization using the AM1^42^ Hamiltonian in the semi-empirical program MOPAC.^35^ The AM1 calculated heats of formation were then used to identify Boltzmann weights for *H*_*shape*_ values for each conformation. An average *H*_*shape*_ was then calculated for each molecule.

### Entropy from chemical properties

For comparison, we calculated entropies from the chemical properties of molecules using ParaSurf. For isodensity surfaces, ParaSurf calculates gas-phase entropy as:

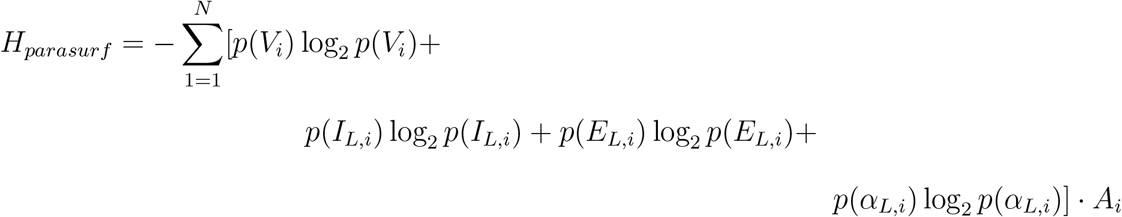

where *p*(*X*_*i*_) is the probability of value *X*_*i*_ property, *n* is the number of triangles on the surface, *A*_*i*_ the area of triangle *i*, and *V*_*i*_, *I*_*L,i*_, *E*_*L,i*_, and *α*_*L,i*_ are the average values of molecular electrostatic potential (*V*), ionization energy (*I*_*L*_), electron affinity (*E*_*L*_), and polarizability (*α*_*L*_), respectively, for triangle *i*.

## Results

We curated experimental gas-phase entropy values of molecules from the literature. In our literature search, most of the molecules happened to be organic compounds, i.e. involving elements C,H,N,O,S,P,Cl,Br, and I. Subsequently, we chose only organic molecules, 1212 in total, for further evaluation. We calculated their structures, charges, molecular surfaces, and shape entropies from their SMILES strings. Please see the Methods section for details of each of the steps. In subsequent sections, we show that the shape entropy has a collinear relationship with the gas-phase entropy. The inclusion of multiple conformations of molecules under study did not improve the *R*^2^ value significantly. From the collinearity, we can estimate the gas-phase of a molecule in a matter of seconds compared to tens to hundreds of CPU hours for other methods. We show that SHAPE, our shape-based entropy method, capture the vibrational entropy of the molecule well. The inclusion of gas-phase entropy in SMINA docking scores improves its performance in predicting binding affinity.

### Gas-phase entropy varies linearly with shape entropy

In our data set, the shape entropy, *H*_*shape*_, showed a linear relationship with gas-phase entropy, *S*_*gasphase*_, with *R*^2^ values of ≈0.9 (Table 2). For calculation of the shape entropy, two different definitions of surfaces were used, (i) the Gaussian surface calculated from the VDW radii of atoms, and (ii) surface calculated from the isodensity of electron of the molecules. Isodensity surfaces of electrons were calculated by including all the atoms of the molecules. For VDW surfaces, we calculated the surfaces by (i) including all the atoms and (ii) excluding the hydrogen atoms. The exclusion of hydrogen atoms improved the *R*^2^ values from 0.83 & 0.84 to 0.91 & 0.88 for 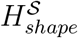 and 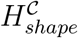, respectively (Table 2). For the remainder of the article, we calculated the VDW surfaces by excluding hydrogens. The linearity of 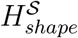 calculated from the VDW surface and 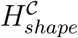 calculated from the isodensity of electron are shown in Figure 1(A) and 1(B), respectively.

### The inclusion of multiple conformations did not improve the collinearity between shape and gas-phase entropy

For calculating *H*_*shape*_ we generated only a single conformation of a molecule. To investigate the possible effect of molecular flexibility on the linear relationship between *H*_*shape*_ and *S*_*gasphase*_, we plotted RMSE values of the linear fit as a function of the number of rotatable bonds (*N*_*rot*_). The RMSE values increase with the *N*_*rot*_, indicating possible dependence of molecular flexibility on the RMSE of fitting (Figure 1(F)). Note that in our curated data, we have *N*_*rot*_ values from 0 to 15; however, in the figure we only plotted values for *N*_*rot*_ 0 to 8, as there are only twelve molecules with values *N*_*rot*_ 9 to 15, an average of 2 molecules for each of those *N*_*rot*_ values. To take into account the effect of flexibility, we

1. generated multiple conformations of each molecule,
2. averaged the *H*_*shape*_ values according to their Boltzmann weights to calculate average *H*_*shape*_, and
3. recalculated the equation of linearity between *H*_*shape*_ and *S*_*gasphase*_, RMSE and *R*^2^values.

The inclusion of multiple conformations did not improve the fit between *H*_*shape*_ with *S*_*gasphase*_ statistically (rows 4 & 5 versus 6 & 7 of the Table 2).

### Comparison between shape entropy and other methods

Gas-phase entropy can be measured at different levels of accuracies with different computational techniques. Generally, the gas-phase entropy is divided into three parts.

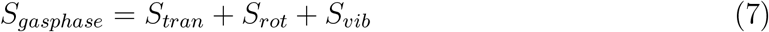

where *S*_*tran*_, *S*_*rot*_, and *S*_*vib*_ are the translational, rotational and vibrational entropies, respectively. Conformational entropy is often considered as an additional part of the gas-phase entropy. The effect of the conformational entropy can be included in the calculation with proper averaging of *S*_*tran*_, *S*_*rot*_, and *S*_*vib*_ arising from different conformations of a molecule and has not been explicitly treated while comparing with other methods. Moreover, conformational entropy accounts for less than 5% of gas-phase entropy in small molecules.^29^ The rotational and translation entropies of a molecule can be calculated from the geometry of the molecule. For vibrational entropy, different levels of theories with different computational resource requirements are used. Out of the 1212 molecules studied, we curated 1091 vibrational entropy values calculated using the quantum-chemical Gaussian-4 (G4) level of theory (Figure 1(C)), a very accurate theory to predict molecular properties, and 389 values calculated using normal mode analysis (NMA) (Figure 1(D)) (see Methods).^27^ The remaining terms of the gas-phase entropy, translational and rotational entropies, were calculated from the shapes of the molecules using analytical methods.^27^ Three different molecular mechanics force-fields, general AMBER force field (GAFF) combined with charges determined by either restrained electrostatic potential fitting (GAFF-ESP) or the AM1-BCC method with bond charge corrections (GAFF-BCC) and the CHARMM general force field (CGenFF) were used for the normal mode calculations.

Three different metrics were used to evaluate the performance of predicting *S*_*gasphase*_ values: i) RMSE, ii) *R*^2^, and iii) mean absolute percentage error (MAPE) from the linear fit with the experimental values. 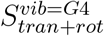 performs best in predicting the gas-phase entropies (row 6 of the Table 2). 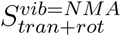 also perform well as shown in row 7 of the Table 2. Readers should note that the slope of the fit for G4 and NMA is not exactly 1. This may be attributed to the fact that the vibration spectra were calculated from a single conformation. Please see the Methods section for further details. As the number of molecules with 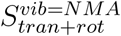 data is only 389, compared to more than 1000 for other methods, the confidence interval is wider for 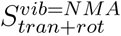. Both the above methods take a considerable amount of time and resources for calculation or parameterization. In comparison, MAPE values of SHAPE are only 1-2% higher, which corresponds to about 0.6 Kcal/mol energy at room temperature, but can be calculated at a fraction of time and resource and without the time and resource-intensive parameterization for every molecule (Table 3). For example, a typical G4 calculation requires hundreds of CPU hours. NMA calculations are fast and can be completed in seconds; however, every new molecule requires new molecular mechanics force field parameterization, taking tens to hundreds of CPU hours. In comparison, SHAPE takes seconds to calculate entropy and does not need any additional parameterization for new molecules. There are two major steps in shape-based entropy calculation, 1) calculation of surface, which has an upper limit runtime complexity of *O*(*logM*), and 2) calculation of histogram, which has an upper limit runtime complexity of *O*(*M*), where *M* is the number of points on the surface. As the number of atoms *N*_*a*_ increases, at worst, the surface area increases linearly with *N*_*a*_. Consequently, the worst-case runtime complexity for shape-entropy is *O*(*N*_*a*_), compared to the best case quadratic runtime complexity 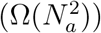 for NMA calculations. For G4 and other quantum mechanical calculations, the runtime complexity is 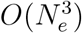 for isolated atoms. It can approach an upper limit of 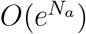 as the number of interacting atoms increases, where *N*_*e*_ is the number of electrons. In our experience, the runtime complexity for G4 calculations has been cubic to the number of atoms 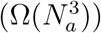. In comparison, in our data, the shape-based entropy exhibited a runtime complexity of 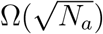. In our dataset, RMSE or MAPE was the lowest for CGenFF in the NMA category. Results for GAFF-ESP and GAFF-BCC are shown in Supporting Information (SI).

### Inclusion of gas-phase entropy may improve binding affinity prediction

To investigate whether the inclusion of gas-phase entropy can improve docking scores, we selected a congeneric series of D-phenyl-proline-based 22 thrombin inhibitors for which experimental binding affinities are available.^43^ The selected ligands were aligned with the co-crystallized ligand (PDB ID: 2zff) using the flexible alignment option in the LS-Align software.^44^ For each aligned ligand, a set of interaction terms were calculated using SMINA:^26^ a Gaussian steric and repulsion term, an electrostatic term, a hydrophobic term, and a hydrogen bond term. We used stepwise (both forward and backward selection) regression^45^ to fit coefficients to these terms. Model performance assessments based *R*^2^, RMSE, and rank correlation coefficient Kendall’s *τ* between the predicted and observed affinities with and without the entropy term are showed in Table 4.

This exercise aims not to draw a statistically robust conclusion, which is beyond the scope of this article and not possible given the small number of data points, rather the goal is to observe the trend as a proof of concept. Consequently, further discussion in this section is based on the mean values only.

The inclusion of gas-phase entropy improved both *R*^2^ and Kendall’s *τ* values from ≈0.20 to 0.40, i.e., from no association between the docking score and binding affinity to a weak association between the two. In this exercise, we have included the gas-phase entropy of the ligand to the docking score, whereas, ideally, one should include the difference in gas-phase entropy between the bound and free form of the proteins and ligands. In our dataset, since all the ligands bind to the same protein pocket, as a first approximation, differences in binding entropy between protein-ligand complexes can be approximated by the differences between the gas-phase entropy of bound ligand conformations. We expect that a proper inclusion of all entropy terms in the docking score will further improve binding affinity prediction.

## Discussion

Calculating entropy is fundamental in calculating the chemical properties of a molecule. The computational resources required to count all possible microstates and their probabilities increase exponentially with the number of atoms. Consequently, other than for very few simple molecules, a direct calculation of entropy becomes infeasible. Typically molecules are approximated as simple harmonic oscillators to calculate entropy. This article has shown an empirical linear relation between shape entropy and gas-phase entropy of small molecules. The empirical relationship estimates gas-phase entropy with 1 − 2% higher error but using 3-4 orders of magnitude fewer resources. Calculation of the gas-phase entropy using SHAPE requires ≈0.01 CPU hours, and its run time complexity is 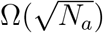, where *N*_*a*_ is the number of atoms. In comparison, a quantum mechanical calculation of vibrational entropy requires ≈100 CPU hours, and the run time complexity of such calculations can approach 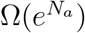. Molecular mechanics-based normal mode analysis takes ≈10 CPU hours for parameterization and the run time complexity is 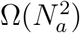. The fast calculation by SHAPE makes it feasible to estimate gas-phase entropy in a demanding computational experiment, e.g., high throughput docking experiments, where millions or billions of such calculations are carried out. The inclusion of gas-phase entropy of ligand in the docking score improved the rank correlation between binding affinity and binding score from 0.18 to 0.40 in our data set.

In our formulation, shape entropy of an isolated atom is ≈0 (Table 1). Consequently, shape entropy is sensitive to the features at the junction of atoms. The accuracy of SHAPE thus depends on the accurate surface description of such atomic junctions. Our calculation observed the increase of collinearity between shape entropy and gas-phase entropy when the VDW surface was calculated without hydrogen atoms (Table 2). We think the VDW surface contour at a junction of hydrogen, the smallest atom, and another atom is most prone to artifacts due to the assumptions in SHAPE. Consequently, removing hydrogen atoms reduces the error of estimation of gas-phase entropy using SHAPE.

**Table 1:**
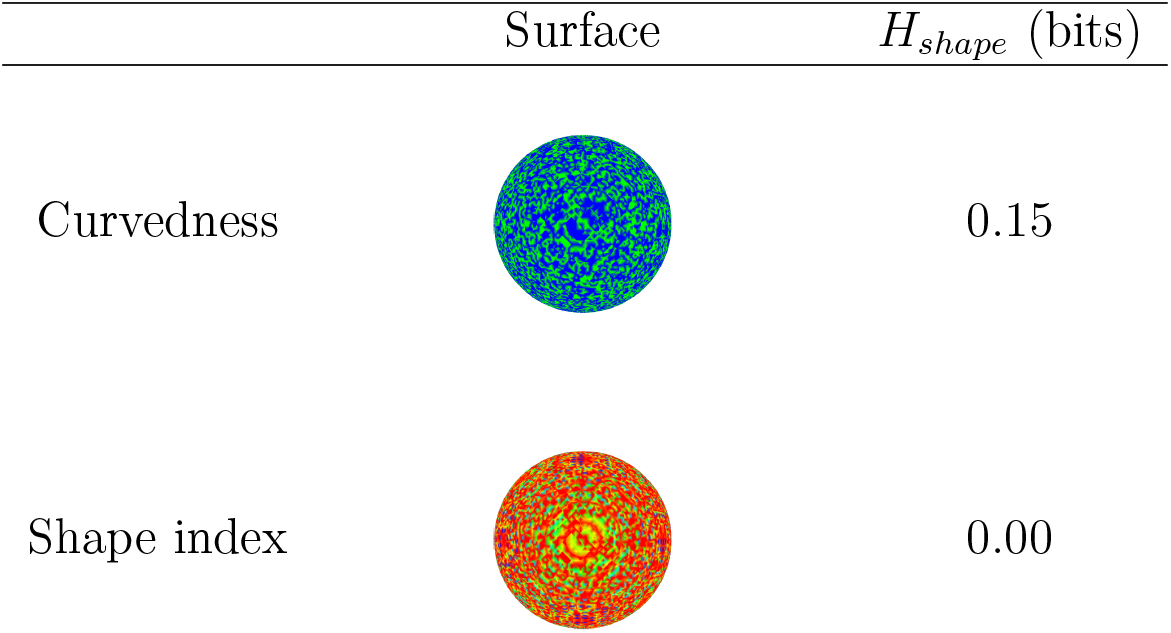
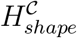 and 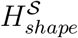 values of a spherical carbon atom. The surface color shading follows a hot cold color scale 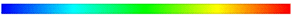 with the low/negative regions coded blue, high/positive regions coloured red and the mid ranges coloured green/yellow.

| | Surface | $H_{shape}$ (bits) |
| --- | --- | --- |
| Curvedness |  | 0.15 |
| Shape index |  | 0.00 |

**Table 2:** The first five rows of the table show the linearity of gas-phase entropy with shape entropy calculated from the VDW surface with and without hydrogen and the isosurface of the electron density with hydrogen. For the VDW surface, we calculated shape entropy from both curvedness 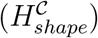 and shape index 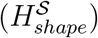. For the isodensity of electrons, we calculated the shape entropy from curvedness. For rows six and seven, multiple conformations were generated for each molecule, and their *H*_*shape*_ values without hydrogen were averaged with Boltzmann weights. Rows eight and nine show the root mean square error (RMSE) and mean absolute percentage error (MAPE) in predicting vibrational entropy by quantum-chemical Gaussian-4 (G4) level of theory and normal mode analysis (NMA) using CHARMM general molecular mechanics force field (CGenFF), respectively, and rotational and translational entropy (*H*_*tran*+*rot*_) analytically form the shape of the molecules (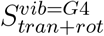 and 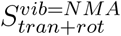). The last row shows the RMSE in fitting the gas-phase entropy values calculated by Parasurf (*H*_*parasurf*_). We used bootstrap methods to calculate lower and upper limits of 95% confidence interval. Upper and lower limits are shown as raised and lowered numbers, respectively.

| Method | $S_{\text{gasphase}} =$<br>($J/mol \cdot K$) | RMSE<br>($J/mol \cdot K$) | R<sup>2</sup> | MAPE<br>% |
| --- | --- | --- | --- | --- |
| Iso with $H_2$ | $139.78_{134.10}^{145.50} + 12.60_{12.28}^{12.92} \times H_{shape}^C + \epsilon$ | $30.05_{28.68}^{31.92}$ | $0.88_{0.86}^{0.90}$ | 6.18 |
| VDW with $H_2$ | $107.72_{100.00}^{116.90} + 9.28_{8.93}^{9.56} \times H_{shape}^S + \epsilon$ | $36.08_{35.52}^{37.89}$ | $0.83_{0.80}^{0.85}$ | 7.56 |
| VDW with $H_2$ | $138.19_{131.20}^{145.30} + 7.03_{6.80}^{7.28} \times H_{shape}^C + \epsilon$ | $35.08_{33.54}^{36.83}$ | $0.84_{0.81}^{0.86}$ | 6.97 |
| VDW | $118.11_{113.50}^{122.90} + 7.73_{7.58}^{7.88} \times H_{shape}^S + \epsilon$ | $25.85_{24.80}^{27.09}$ | $0.91_{0.90}^{0.92}$ | 5.35 |
| VDW | $117.50_{111.30}^{123.70} + 6.68_{6.50}^{6.85} \times H_{shape}^C + \epsilon$ | $29.70_{28.39}^{31.26}$ | $0.88_{0.86}^{0.90}$ | 5.90 |
| VDW with BW | $121.55_{116.60}^{126.50} + 7.60_{7.43}^{7.76} \times H_{shape}^S + \epsilon$ | $26.41_{25.35}^{27.64}$ | $0.91_{0.89}^{0.92}$ | 5.41 |
| VDW with BW | $122.07_{117.00}^{127.10} + 6.48_{6.34}^{6.63} \times H_{shape}^C + \epsilon$ | $28.29_{27.10}^{29.64}$ | $0.89_{0.88}^{0.91}$ | 5.66 |
| G4+analytical | $-48.33_{-54.08}^{-42.48} + 1.17_{1.15}^{1.19} \times S_{tran+rot}^{vib=G4} + \epsilon$ | $16.35_{15.52}^{17.37}$ | $0.96_{0.95}^{0.96}$ | 3.33 |
| NMA+analytical | $-34.79_{-52.95}^{-13.43} + 1.11_{1.05}^{1.17} \times S_{tran+rot}^{vib=NMA} + \epsilon$ | $22.45_{15.55}^{27.08}$ | $0.88_{0.82}^{0.92}$ | 4.15 |
| Parasurf | $170.42_{157.60}^{182.20} + 6.23_{5.84}^{6.65} \times H_{parasurf} + \epsilon$ | $59.45_{55.65}^{64.33}$ | $0.52_{0.48}^{0.57}$ | 11.05 |

**Table 3:** A comparison of resources needed for calculating gas-phase entropy and parameterization for the calculation, the runtime complexity, and mean average percentage error (MAPE) for different methods. Estimates of resources are for molecules of similar sizes in our database. *N*_*a*_ is the number of atoms. For G4 and other quantum mechanical calculations, the runtime complexity is at least cubic 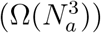. See the text for more details. For NMA calculations, ≈10 CPU hours are needed for the parameterization of each ligand.

| Method | Parameterization<br>CPU hours | Calculation<br>CPU hours | Runtime<br>Complexity | Error<br>(MAPE) | Hi-throughput<br>docking |
| --- | --- | --- | --- | --- | --- |
| G4+analytical | **0** | $\approx 100$ | $\Omega(N_a^3)$ | **3.33%** | **×** |
| NMA+analytical | $\approx 10$ | $\approx \mathbf{0.01}$ | $\Omega(N_a^2)$ | 4.15% | **×** |
| **Shape** | **0** | $\approx \mathbf{0.01}$ | $\Omega(\sqrt{N_a})$ | 5.35% | **✓** |

**Table 4:** Performance of the scoring functions for determining binding affinity of thrombin ligands with and without including the shape entropy term.

|  | **RMSE**<br>(Kcal/mol) | **R<sup>2</sup>** | **Kendall’s $\tau$** |
| --- | --- | --- | --- |
| $-H_{shape}$ | $1.35^{1.00}_{1.92}$ | $0.23^{0.71}_{0.01}$ | $0.18^{0.47}_{-0.10}$ |
| $+H_{shape}$ | $1.18^{1.00}_{1.61}$ | $0.40^{0.61}_{0.03}$ | $0.40^{0.55}_{-0.23}$ |

Shape entropy has previously been used to identify different chemical properties, such as the ionization potential. The underlying relationship between shape entropy and various chemical and thermodynamical properties is still an active research area. Further studies are needed to investigate the underlying relationship between molecular shape, shape entropy, and molecular properties. This article shows the power of shape entropy in estimating thermodynamical properties.

## Conclusion

Shape entropy is a promising tool to estimate the thermodynamical properties of molecules. We have shown that shape entropy can be used for a fast and accurate estimation of gas-phase entropy. This opens up the possibility of including a direct estimate of entropy in scoring functions for docking, where fast calculations are needed in high-throughput docking. We have further shown that the inclusion of ligand gas-phase entropy improves docking results. For a proper treatment, solvent and protein entropy also need to be considered, which are work in progress.

## Acknowledgement

VV acknowledges financial support from the Research Council of Norway (Grant No. 262152). AR acknowledges funding from the National Institute of Allergy and Infectious Diseases (NIAID), National Institutes of Health (NIH), Department of Health and Human Services under BCBB Support Services Contract HHSN316201300006W/HHSN27200002 to MSC, Inc. The authors thank Cepos InSilico Ltd. for providing an academic license for ParaSurf. The authors thank Drs Travis Wheeler and Natalia Syzochenko for their valuable input about the manuscript. The authors thank Ryan Kissinger of Visual and Medical Arts, Research and Technology Branch, NIAID, NIH for his valuable input in the graphical table of content.

## Supporting Information Available

1. An excel file with the observed and predicted gas-phase entropies for the different compounds (in SMILES format), including gas-phase entropy calculated using NMA with the molecular mechanics force fields GAFF-ESP and GAFF-BCC.
2. A table showing the effect of smoothing parameter *σ* on the surface calculation.
3. A figure showing the effects of smoothing parameter *σ* and number of bins for calculating histogram on the shape entropy.

## References

(1) Parks, C. D.; Gaieb, Z.; Chiu, M.; Yang, H.; Shao, C.; Walters, W. P.; Jansen, J. M.; McGaughey, G.; Lewis, R. A.; Bembenek, S. D.; et al. D3R grand challenge 4: blind prediction of protein–ligand poses, affinity rankings, and relative binding free energies. J. Comput. Aided Mol. Des. 2020, 34, 99–119.

(2) Gaieb, Z.; Parks, C. D.; Chiu, M.; Yang, H.; Shao, C.; Walters, W. P.; Lambert, M. H.; Nevins, N.; Bembenek, S. D.; Ameriks, M. K.; et al. D3R Grand Challenge 3: blind prediction of protein–ligand poses and affinity rankings. J. Comput. Aided Mol. Des. 2019, 33, 1–18.

(3) Guedes, I. A.; Pereira, F. S.; Dardenne, L. E. Empirical scoring functions for structure-based virtual screening: applications, critical aspects, and challenges. Front. Pharmacol. 2018, 9, 1089.

(4) Winkler, D. A. Ligand Entropy Is Hard but Should Not Be Ignored. J. Chem. Inf. Model. 2020, 60, 4421–4423.

(5) Gilson, M. K.; Zhou, H.-X. Calculation of protein-ligand binding affinities. Annu. Rev. Biophys. Biomol. Struct. 2007, 36.

(6) Chia-en, A. C.; Chen, W.; Gilson, M. K. Ligand configurational entropy and protein binding. Proc. Natl. Acad. Sci U. S. A 2007, 104, 1534–1539.

(7) Lill, M. A. Efficient incorporation of protein flexibility and dynamics into molecular docking simulations. Biochemistry 2011, 50, 6157–6169.

(8) Niessen, K. A.; Xu, M.; Paciaroni, A.; Orecchini, A.; Snell, E. H.; Markelz, A. G. Moving in the right direction: protein vibrations steering function. Biophys. J. 2017, 112, 933–942.

(9) Cooper, A.; Dryden, D. Allostery without conformational change. Eur. Biophys. J. 1984, 11, 103–109.

(10) Shannon, C. E. A mathematical theory of communication. Bell Syst. Tech. J. 1948, 27, 379–423.

(11) Gao, X.; Gallicchio, E.; Roitberg, A. E. The generalized Boltzmann distribution is the only distribution in which the Gibbs-Shannon entropy equals the thermodynamic entropy. J. Chem. Phys. 2019, 151, 034113.

(12) Rong, C.; Wang, B.; Zhao, D.; Liu, S. Information-theoretic approach in density functional theory and its recent applications to chemical problems. Wiley Interdiscip. Rev. Comput. Mol. Sci. 2019, e1461.

(13) Pineda-Urbina, K.; Guerrero, R.; Reyes, A.; Gómez-Sandoval, Z.; Flores-Moreno, R. Shape entropy’s response to molecular ionization. J. Mol. Model. 2013, 19, 1677–1683.

(14) Guthrie, J. P. Use of DFT Methods for the Calculation of the Entropy of Gas Phase Organic Molecules: An Examination of the Quality of Results from a Simple Approach. J. Phys. Chem. A 2001, 105, 8495–8499.

(15) Rozanska, X.; Stewart, J. J. P.; Ungerer, P.; Leblanc, B.; Freeman, C.; Saxe, P.; Wimmer, E. High-Throughput Calculations of Molecular Properties in the MedeA Environment: Accuracy of PM7 in Predicting Vibrational Frequencies, Ideal Gas Entropies, Heat Capacities, and Gibbs Free Energies of Organic Molecules. J. Chem. Eng. Data 2014, 59, 3136–3143.

(16) Ghahremanpour, M. M.; van Maaren, P. J.; Ditz, J. C.; Lindh, R.; van der Spoel, D. Large-scale calculations of gas phase thermochemistry: Enthalpy of formation, standard entropy, and heat capacity. J. Chem. Phys. 2016, 145, 114305.

(17) Vansteenkiste, P.; Speybroeck, V. V.; Marin, G. B.; Waroquier, M. Ab Initio Calculation of Entropy and Heat Capacity of Gas-Phasen-Alkanes Using Internal Rotations. J. Phys. Chem. A 2003, 107, 3139–3145.

(18) Červinka, C.; Fulem, M.; Růžička, K. Evaluation of Uncertainty of Ideal-Gas Entropy and Heat Capacity Calculations by Density Functional Theory (DFT) for Molecules Containing Symmetrical Internal Rotors. J. Chem. Eng. Data 2013, 58, 1382–1390.

(19) Barrett, R. A.; Meier, R. J. The calculation of molecular entropy using the semiempirical AM1 method. J. Mol. Struct. THEOCHEM 1996, 363, 203–209.

(20) Åqvist, J.; Medina, C.; Samuelsson, J.-E. A new method for predicting binding affinity in computer-aided drug design. Protein Eng Des Sel. 1994, 7, 385–391.

(21) Gouda, H.; Kuntz, I. D.; Case, D. A.; Kollman, P. A. Free energy calculations for theophylline binding to an RNA aptamer: comparison of MM-PBSA and thermodynamic integration methods. Biopolymers: Original Research on Biomolecules 2003, 68, 16–34.

(22) Straatsma, T.; McCammon, J. Multiconfiguration thermodynamic integration. J. Chem. Phys. 1991, 95, 1175–1188.

(23) Domalski, E. S.; Hearing, E. D. Estimation of the Thermodynamic Properties of Hydrocarbons at 298.15 K. J. Phys. Chem. Ref. Data 1988, 17, 1637–1678.

(24) Sabbe, M. K.; Vleeschouwer, F. D.; Reyniers, M.-F.; Waroquier, M.; Marin, G. B. First Principles Based Group Additive Values for the Gas Phase Standard Entropy and Heat Capacity of Hydrocarbons and Hydrocarbon Radicals. J. Phys. Chem. A 2008, 112, 12235–12251.

(25) Blurock, E. S.; Warth, V.; Grandmougin, X.; Bounaceur, R.; Glaude, P.-A.; Battin-Leclerc, F. JTHERGAS: Thermodynamic estimation from 2D graphical representations of molecules. Energy 2012, 43, 161 – 171.

(26) Koes, D. R.; Baumgartner, M. P.; Camacho, C. J. Lessons Learned in Empirical Scoring with smina from the CSAR 2011 Benchmarking Exercise. J. Chem. Inf. Model. 2013, 53, 1893–1904.

(27) van der Spoel, D.; Ghahremanpour, M. M.; Lemkul, J. A. Small Molecule Thermo-chemistry: A Tool for Empirical Force Field Development. J. Phys. Chem. A 2018, 122, 8982–8988.

(28) Raychaudhury, C.; Rizvi, I.; Pal, D. Predicting gas phase entropy of select hydrocarbon classes through specific information-theoretical molecular descriptors. SAR QSAR Environ. Res. 2019, 30, 491–505.

(29) Chan, L.; Morris, G.; Hutchison, G. Understanding Conformational Entropy in Small Molecules. 2020,

(30) Landrum, G. RDKit: Open-Source Cheminformatics Software. 2020; RDKit version 2020.09.1.0.

(31) O’Boyle, N. M.; Banck, M.; James, C. A.; Morley, C.; Vandermeersch, T.; Hutchison, G. R. Open Babel: An open chemical toolbox. J Cheminf. 2011, 3.

(32) Rappe, A. K.; Casewit, C. J.; Colwell, K. S.; Goddard, W. A.; Skiff, W. M. UFF, a full periodic table force field for molecular mechanics and molecular dynamics simulations. J Am. Chem. Soc. 1992, 114, 10024–10035.

(33) Gabdoulline, R.; Wade, R. Analytically defined surfaces to analyze molecular interaction properties. J. Mol. Graph. 1996, 14, 341–353.

(34) Clark, T. ParaSurf 12. 2012; Cepos InSilico Ltd.

(35) Stewart, J. J. P. MOPAC2016. 2016; Stewart Computational Chemistry, Colorado Springs, CO, USA, (http://OpenMOPAC.net).

(36) do Carmo, M. P. Differential geometry of curves and surfaces., Prentice Hall, 1976; pp I–VIII, 1–503.

(37) Goldman, R. Curvature formulas for implicit curves and surfaces. Comput. Aided Geom. Des. 2005, 22, 632–658, Geometric Modelling and Differential Geometry.

(38) Xia, K.; Feng, X.; Chen, Z.; Tong, Y.; Wei, G.-W. Multiscale geometric modeling of macromolecules I: Cartesian representation. J. Comput. Phys. 2014, 257, 912–936.

(39) Koenderink, J. J.; van Doorn, A. J. Surface shape and curvature scales. Image Vis Comput 1992, 10, 557–564.

(40) Schlager, S. In Statistical Shape and Deformation Analysis; Zheng, G.; Li, S.; Szekely, G., Eds., Academic Press, 2017; pp 217–256.

(41) Popinet, S. GTS: GNU Triangulated Surface Library. http://gts.sourceforge.net/, 2010.

(42) Dewar, M. J. S.; Zoebisch, E. G.; Healy, E. F.; Stewart, J. J. P. Development and use of quantum mechanical molecular models. 76. AM1: a new general purpose quantum mechanical molecular model. J Am. Chem. Soc. 1985, 107, 3902–3909.

(43) Baum, B.; Mohamed, M.; Zayed, M.; Gerlach, C.; Heine, A.; Hangauer, D.; Klebe, G. More than a Simple Lipophilic Contact: A Detailed Thermodynamic Analysis of Non-basic Residues in the S1 Pocket of Thrombin. J. Mol. Biol. 2009, 390, 56–69.

(44) Hu, J.; Liu, Z.; Yu, D.-J.; Zhang, Y. LS-align: an atom-level, flexible ligand structural alignment algorithm for high-throughput virtual screening. Bioinformatics 2018, 34, 2209–2218.

(45) Zhang, Z. Variable selection with stepwise and best subset approaches. Ann. Transl. Med. 2016, 4, 136–136.

