## Supplementary figures and Data for "Fast and Accurate Estimation of Gas-Phase Entropy from the Molecular Surface Curvature": Venkatraman_Roy_SI.pdf

### Supporting Information for Fast and Accurate Estimation of Gas-Phase Entropy from the Shape of a Molecule

E-mail:

*Table ST1: Effect of the smoothing factor $\sigma$ on the molecular surface. Larger values of $\sigma$ smooth out the details of the surface, while the smaller values of $\sigma$ preserve more details of the surface features. In this article $\sigma = 0.1$ has been used. Please see the main text for definitions of Curvedness and Shape Index. The colour scheme is based on the standard blue to green to red colour map where low values of the property are shown in blue, values close to the maxima in red and those in between in green.*

| $\sigma$ | *Curvedness* | *Shape Index* |
|---|---|---|
| 0.1 | 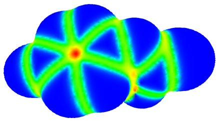 | 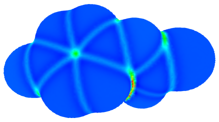 |
| 0.3 | 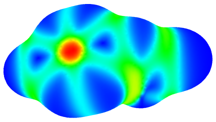 | 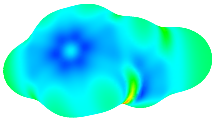 |
| 0.5 | 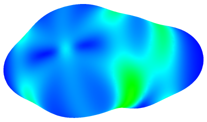 | 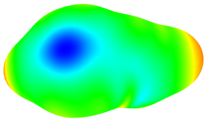 |

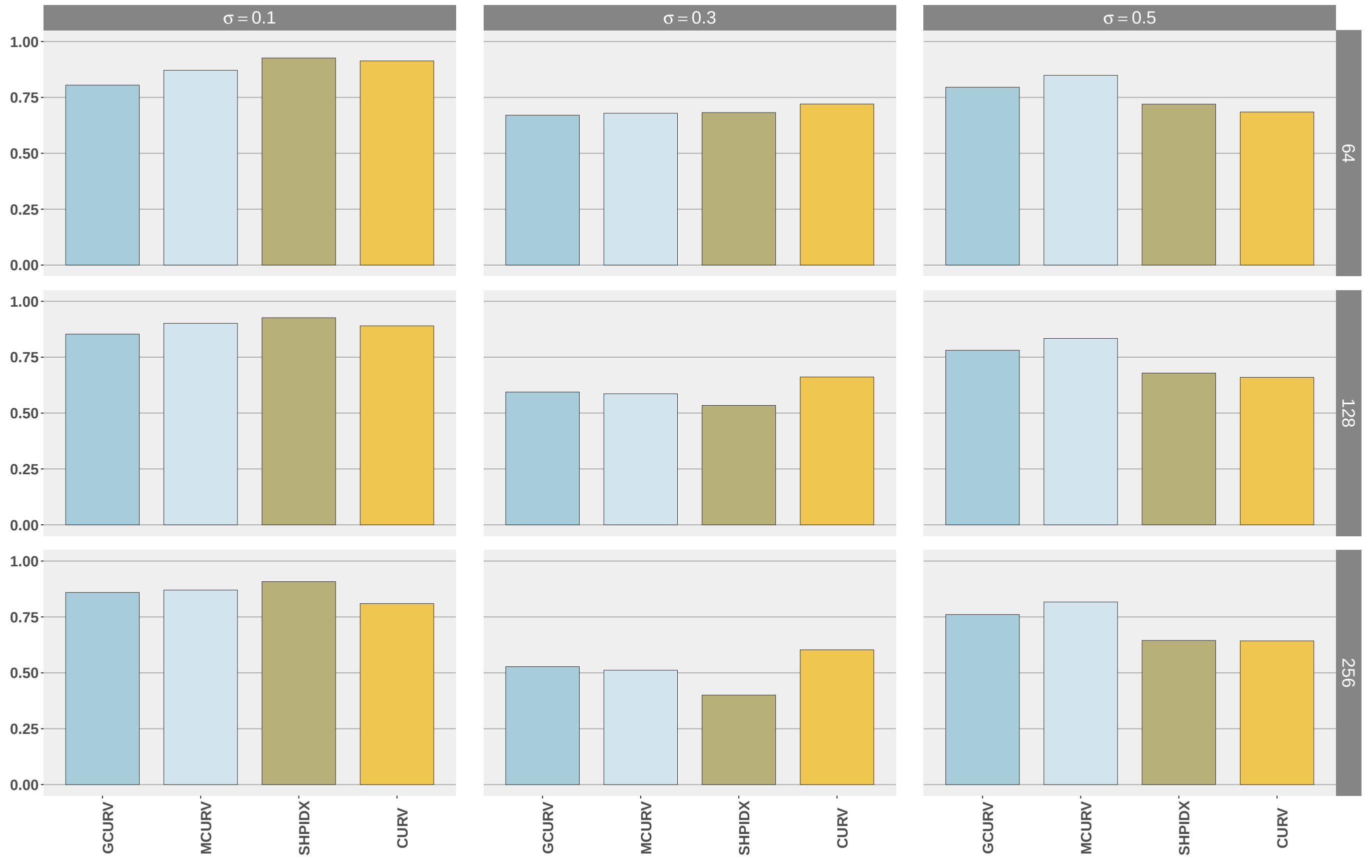


Figure SF1: The figure shows the Pearson correlation coefficients between experimental gas-phase and the shape entropy values calculated using different definitions of surface curvature.The shape Index (SHPIDX) and curvedness (CURV) are defined in the main text.The Gaussian curvature (GCURV) is defined as $\kappa_1 \times \kappa_2$ and mean curvature (MCURV) is defined as $(\kappa_1 + \kappa_2)/2$. $\kappa_1$ and $\kappa_2$ are principal curvatures and defined in the main text. The correlations between the calculated property-based entropies and the experimental values are shown as barplots (y1-scale). Different panels correspond to the different combinations of the smoothing factor $\sigma$ and the number of bins. The y2-scale shows the number of bins used to calculate the histogram of the surface curvatures and the probability distribution of the curvature values. The correlation values have only a weak dependency on the number of bins. The smoothing factor $\sigma$ has a stronger effect on correlations, which is to be expected as $\sigma$ controls the details of the surface features. We have used 64 bins to calculate the histogram, $\sigma = 0.1$, and SHPIDX and CURV to calculate the shape entropies in this work.
